# Adsorption of RNA to interfaces of biomolecular condensates enables wetting transitions

**DOI:** 10.1101/2023.01.12.523837

**Authors:** Nadia A. Erkamp, Mina Farag, Daoyuan Qian, Tomas Sneideris, Timothy J. Welsh, Hannes Ausserwöger, David A. Weitz, Rohit V. Pappu, Tuomas P. J. Knowles

## Abstract

Biomolecular condensates form via spontaneous and driven phase transitions of multivalent proteins and nucleic acids. These macromolecules can be organized in spatially inhomogeneous ways that lead to multiple coexisting dense phases with distinct macromolecular interfaces. While considerable attention has focused on the physical driving forces that give rise to phase separation from bulk solutions, the interactions that underlie adsorption driven wetting transitions remain unclear. Here, we report that pyrimidine-rich RNAs function as adsorbents that enable cascades of wetting transitions that include partial and complete wetting of condensates formed by purine-rich RNAs. Computations show that macromolecules that are scaffolds of condensates are oriented perpendicular to condensate interfaces whereas adsorbents are oriented parallel to interfaces. Our results yield heuristics for the design of synthetic materials that can be based on RNA-rich condensates featuring bespoke interfaces and distinct local microenvironments created by the interplay between scaffolds versus adsorbents.

---

Biomolecular condensates are compositionally distinct membraneless bodies that form via a combination of spontaneous and driven phase transitions such as phase separation of macromolecules from bulk solutions ^1^. Condensates are thought to provide spatial and temporal control and organization over biochemical reactions involving proteins and nucleic acids, and there is growing interest in developing synthetic condensates with bespoke cellular functions ^2, 3^. These efforts build on our growing understanding of the physico-chemical principles that drive the formation and dissolution of condensates in live cells ^4, 5^, and facsimiles of condensates *in vitro* ^6^. In live cells, condensates comprise several distinct types of macromolecules ^7^. Further, within condensates, one often observes a hierarchical organization of the constituent macromolecules, and this has been reported for stress granules ^8^, nucleoli ^9^, nuclear speckles ^10^, and the mitochondrial nucleoid ^11^. Spatially organized condensates were also shown to arise in the simplest of mixtures^12^ including ternary systems comprising two different RNA molecules and an arginine-rich peptide ^13^.

Long or small non-coding RNA (ncRNA) molecules feature prominently in several nuclear bodies such as nuclear speckles ^10^, peri-nucleolar compartments ^14^, and paraspeckles ^15^. These lncRNA molecules are biological instantiations of flexible, linear polyelectrolytes. In nuclear speckles, the long ncRNA *MALAT1* and small nuclear RNAs localize to the periphery, forming an adsorbed layer on the dense phase that forms around highly repetitive proteins including SON and SC35 ^10^. The interplay of macromolecule-solvent and inter-macromolecule interactions determines the inhomogeneous spatial organization and the compositions of coexisting dense phases within nuclear speckles. In general, some RNA molecules constitute the inner cores of dense phases, whereas others either make up internal interfaces or the main components of interfaces that interact with the condensate on one side and the coexisting dilute phase on the other. In effect, RNA molecules can be scaffolds that drive phase separation from the bulk or they can adsorb to interfaces of condensates formed by scaffolds. *MALAT1* is an example of an RNA that functions as an adsorbent ^15^.

Here, we investigate the different effects of RNA molecules as scaffolds versus adsorbents. Our investigations are motivated by a general framework anchored in the Gibbs adsorption isotherm ^16^, and the physics of adsorption mediated wetting transitions ^17^. We show that the phenomenology from the physics of wetting and adsorption can be realized for model systems comprising a biocompatible polymer, namely polyethylene glycol (PEG) ^18^, and two homopolymeric RNA molecules *viz*., poly-adenine (poly-rA), which is a polypurine, and poly-cytosine (poly-rC), which is a polypyrimidine. We use poly-rA and poly-rC because they are non-base-pairing sequences whose homotypic interactions are likely to be driven by base-stacking. Importantly, they are useful mimics of ncRNA molecules because they cannot form stable secondary and tertiary structures.

## Results

### RNA molecules undergo a concentration-dependent adsorption and wetting transition

Mixtures of poly-rA and poly-rC were combined with PEG to form condensates with a wide variety of mesoscale structures. **Figure 1** shows samples where poly-rA and PEG form condensates. To this two-phase system, we added different amounts of poly-rC. We observed dose-dependent adsorption of poly-rC at the interface between the dilute and dense phases of the pre-formed condensate. Adsorption becomes visible at a mass concentration of 500 ng/μL and is manifest in the form of interfacial foci (**Supplementary Figure 1**). We refer to this as partial wetting ^17^. The number of foci increases in number with increasing concentration of poly-rC (**Supplementary video 1**). Above a threshold concentration that lies between 750 ng/μL and 2000 ng/μL, we observed complete wetting, implying that the system has undergone a wetting transition that leads to two coexisting dense phases. The second dense phase is enriched in poly-rC and it completely wets the interface of the poly-rA-rich condensate. Line profile analysis of the fluorescence intensities showed that poly-rC also adsorbed to the interface between the two dense phases (**Supplementary Figure 2**).

**Figure 1:**
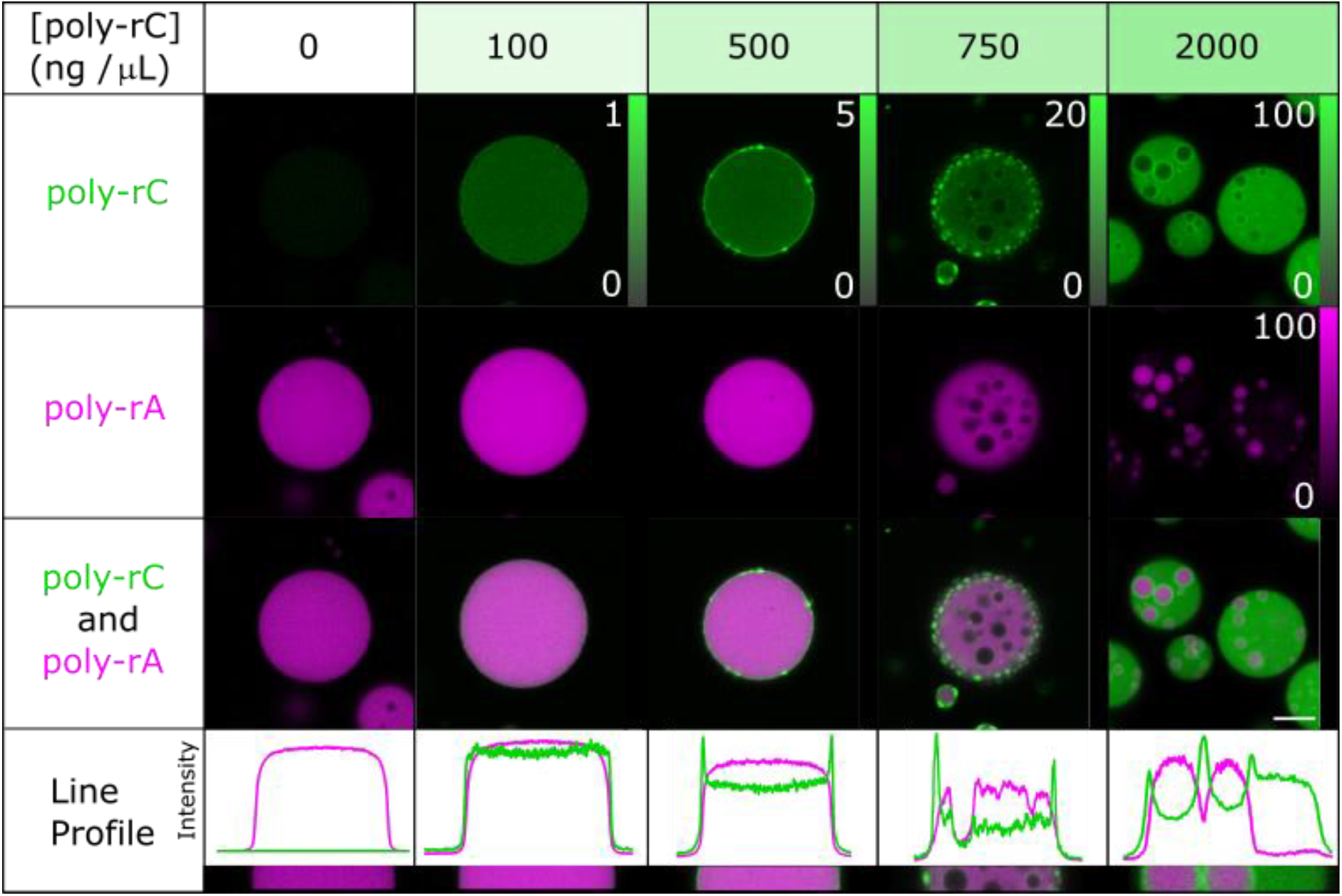
poly-rC adsorbs to and wets the interface of poly-rA-rich condensatesnn. The addition of increased amounts of poly-rC to poly-rA-rich condensates leads to adsorption-mediated transitions from partial to complete wetting ^17^. The wetting transition gives rise to two dense phases, one rich in poly-rA and the other rich in poly-rC. The data suggest adsorption of poly-rC to the interface between the two coexisting dense phases. Samples contain 10^3^ ng/μL poly-rA, 5 w/w% PEG, 750 mM NaCl, and 50 mM HEPES at pH 7.3. The mass concentration of poly-rC is systematically titrated. Legends for relative intensities apply to the images to the left of each legend. The scale bar of 5 μm applies to all images.

### Wetting transitions are governed by relative stoichiometries of poly-rA and poly-rC

PEG-RNA interactions provide the driving forces for forming condensates in the model systems studied in our work ^18, 19^. Increasing the concentration of PEG should alter the chemical potential of PEG, thereby promoting phase separation. Likewise, to zeroth order, increasing the concentration of NaCl should weaken the electrostatic repulsions between RNA molecules, and this should also promote phase separation. As a result, we titrated NaCl and PEG concentrations to tune the overall phase behaviour. The resulting condensates have several features: they can be homogeneous; poly-rC can adsorb to the interface to either partially or completely wet the condensate; or percolated networks can form, especially at high salt concentrations (left to right **Figure 2a**). We also measured coexistence curves of poly-rC by titrations of NaCl and PEG (**Figure 2b**). When the NaCl and PEG concentrations are low, we did not observe phase separation. However, increasing the NaCl and PEG concentrations drives the formation of homogenous condensates (circles).

**Figure 2:**
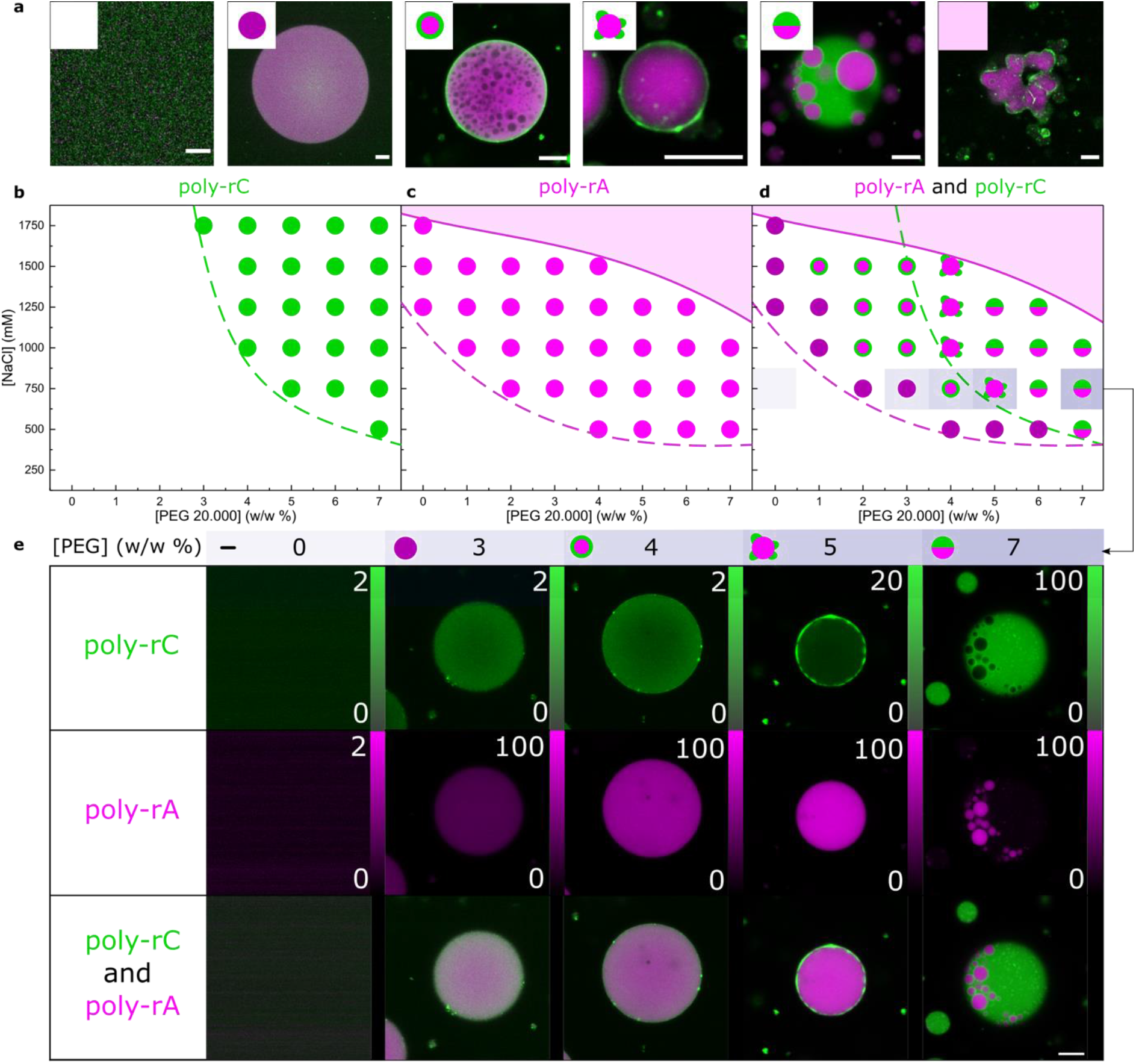
Adsorption and wetting as a function of PEG and NaCl concentrations. **a**. Examples of one-phase, well-mixed solutions, homogeneous poly-rA condensates, adsorption, partial wetting, and complete wetting of poly-rC, and the formation of percolated, gel-like poly-rA structures that are observed at different NaCl and PEG concentrations. Phase boundaries as a function of NaCl and PEG concentration, for **b**. 10^3^ ng/μL poly-rC, **c**. 10^3^ ng/μL poly-rA and **d**. both 10^3^ ng/μL poly-rC and 10^3^ ng/μL poly-rA. Condensates are formed when the phase boundary of poly-rA is crossed, and the type of mesoscale structures that form will depend on the ratio of poly-rC-to-poly-rA. We observe adsorption of poly-rC to the interface of poly-rA condensates in regimes where poly-rC cannot make condensates on its own. **d**. Adding PEG increases the amount of poly-rA in the condensate and causes poly-rC to wet or adsorb onto the condensate in a manner dependent on the relative stoichiometries. Line profiles are shown in Supplementary Figure 3. Samples contain 10^3^ ng/μL poly-rA and/or poly-rC, a variable amount of PEG and NaCl, and 50 mM HEPES pH = 7.3. The scale bar representing 10 μm applies to all confocal images.

We also measured coexistence curves for poly-rA and observed that it forms condensates at lower concentrations of NaCl and PEG than poly-rC (**Figure 2c**). This shows that polypurine systems (poly-rA) drive phase separation through stronger interactions when compared to polypyrimidine systems (poly-rC) ^13, 20^. Strong interactions can engender dynamically arrested phase separation ^13,11^, giving rise to irregular mesoscale structures that are dominated by long-lived intermolecular crosslinks. The resultant percolated networks were observed for poly-rA and not for poly-rC.

Next, we mapped phase boundaries in the presence of equal amounts of poly-rC and poly-rA (**Figure 2d-e**; line profiles in **Supplementary Figure 3**). Condensates were formed in the same concentration regimes of PEG and NaCl as poly-rA by itself. However, the condensates can have various structures. The condensates (purple circle) that are rich in poly-rA formed once the phase boundary of poly-rA is crossed. Upon further increases in NaCl or PEG concentrations, the concentration of poly-rA within these condensates increased. In this regime, poly-rC adsorbs to the interface of the poly-rA-rich condensate, despite being in a regime where it cannot phase separate on its own (green ring around magenta circle). In the concentration regime where the intrinsic phase boundary of poly-rC was crossed, we observed condensates enriched in poly-rC that partially wet the interface of the poly-rA-rich condensate (magenta circle with green droplets on it). Further increases to the concentrations of NaCl or PEG led to the formation of multiphase condensates where a poly-rC-rich dense phase completely wetted a poly-rA-rich dense phase (half magenta, half green circle). These observations show that the relative stoichiometries, specifically PEG:poly-rC and PEG:poly-rA ratios, and the locations with respect to intrinsic phase boundaries determine the adsorption and wetting behaviours of poly-rC.

### poly-rC interacts favourably with interfaces of poly-rA condensates

To explain why poly-rC wets the interface of a condensate, we compared the interaction strengths between the RNA molecules and PEG. We will refer to poly-rA and poly-rC as poly-rX in this section. We used a previously established microfluidics platform^19, 21^ to prepare over 10^5^ samples with varying amounts of poly-rX and PEG (**Figure 3a**). The images were analysed to determine if condensates form, to quantify the total poly-rX and PEG concentrations, and the concentrations of poly-rX in the dilute phase (**Figure 3b**). If the sample did not contain condensates, we labelled this datapoint as being part of the well-mixed one-phase regime (blue). If condensates were observed, we grouped the datapoint depending on the dilute phase poly-rX concentration (red, orange, yellow). Fitting boundaries between these groups of data, we obtained the phase boundary the delineates mixed, and phase separated samples (dashed line). Additionally, the lines fitted between the red, orange, and yellow datasets show the tie-lines (method and materials).

**Figure 3:**
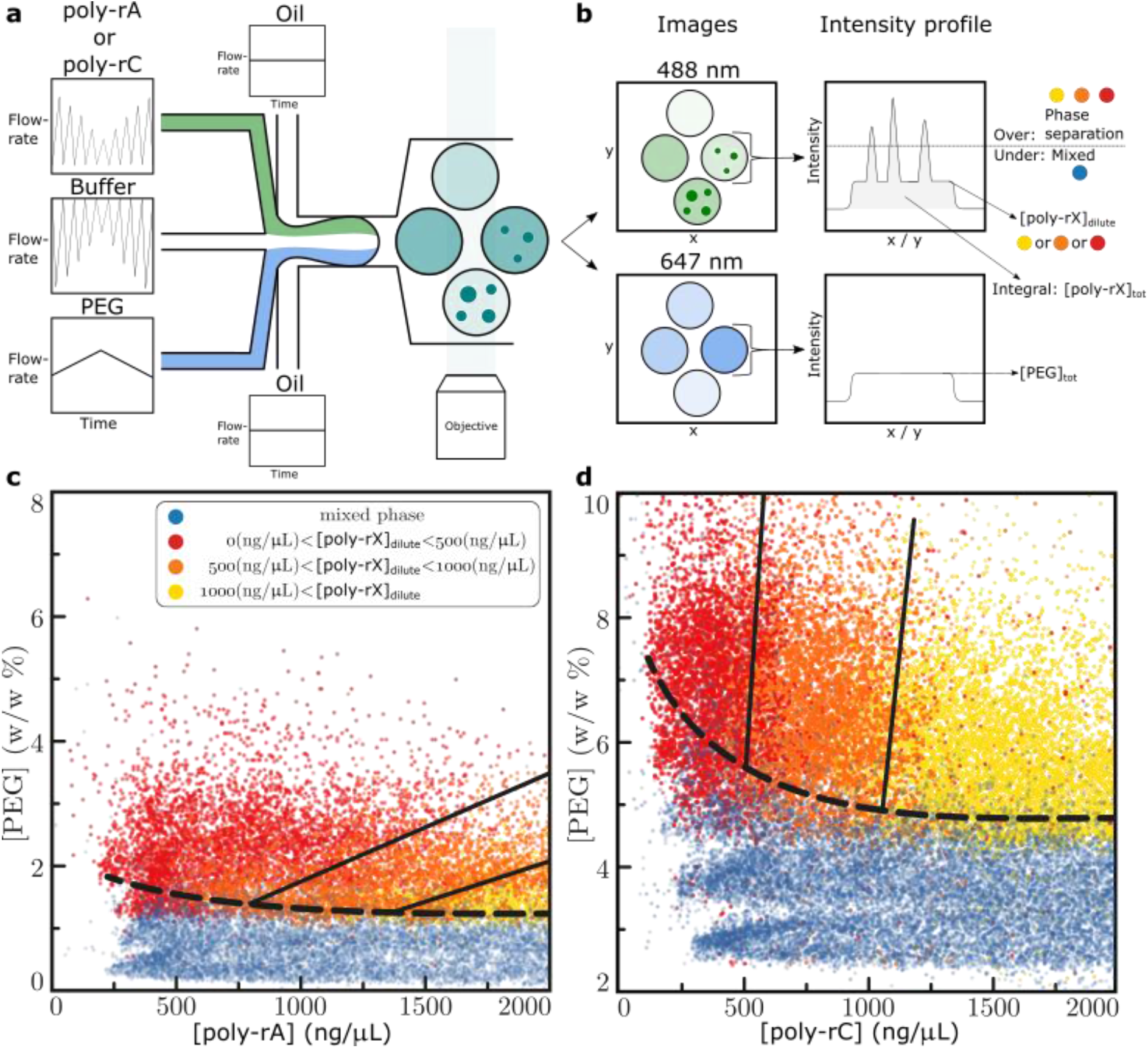
Measurements of phase boundaries using microfluidics. **a**. Schematic describing the microfluidic platform. **b**. Schematic describing how tie-lines are determined. **c-d**. Phase diagrams of poly-rA and poly-rC (poly-rX) generated using a microfluidic platform^21^. Solid lines indicate tie lines, whereas dashed lines indicate phase boundaries. We obtain the tie-lines by grouping data (over 10^5^ datapoints per graph) with condensates based on the dilute phase concentration of poly-rA or poly-rC. The slope of the tie-line for poly-rA is 0.0015 ± 0.0002 and for poly-rC is 0.062 ± 0.016 %/ ng/μL. Molecules that adsorb onto the interface of a condensate have a favourable, but much weaker interaction with the molecules in the condensate, allowing them to adsorb to the interface, rather than being recruited into the condensate (Supplementary Figure 4).

The slopes of tie-lines allow us to quantify the relative contributions of homotypic versus heterotypic interactions in the poly-rX and PEG system. A positive slope indicates that favourable heterotypic interactions drive the formation of condensates, whereas a negative slope indicates that heterotypic interactions are unfavourable. From the data in **Figure 3**, we find tie-lines with slopes of 0.0015 ± 0.0002 and 0.062 ± 0.016 %/ ng/μL for poly-rA and poly-rC, respectively. Thus, both poly-rA and poly-rC interact favourably with PEG ^18, 19^. Notably, as seen in **Figure 2d**, poly-rC can be gradually excluded from the condensate to accommodate more favourable poly-rA-PEG interactions. Since the interactions between poly-rC and PEG are favourable, albeit not as strong as the poly-rA-PEG interactions, the poly-rC molecules adsorb to the interface of the PEG-rich condensate. The weaker interactions between poly-rC and PEG versus poly-rA and PEG are due to the polypurine versus polypyrimidine natures of poly-rA and poly-rC, respectively.

### How is adsorption affected by differences in interaction strengths?

Next, we used a computational coarse-grained model that uses LaSSI ^22^, a lattice-based simulation engine (**Figure 4**), to understand how macromolecules interact preferentially with the interfaces of condensates. We performed simulations involving three distinct homopolymers labelled A, B, and C, each 250 chemical monomers in length. In these models, polymers A, B, and C are designed to mimic poly-rA, PEG, and poly-rC, respectively. Accordingly, polymer B is parameterized to be the scaffold macromolecule that drives condensate formation via heterotypic interactions with polymers A and C.

**Figure 4:**
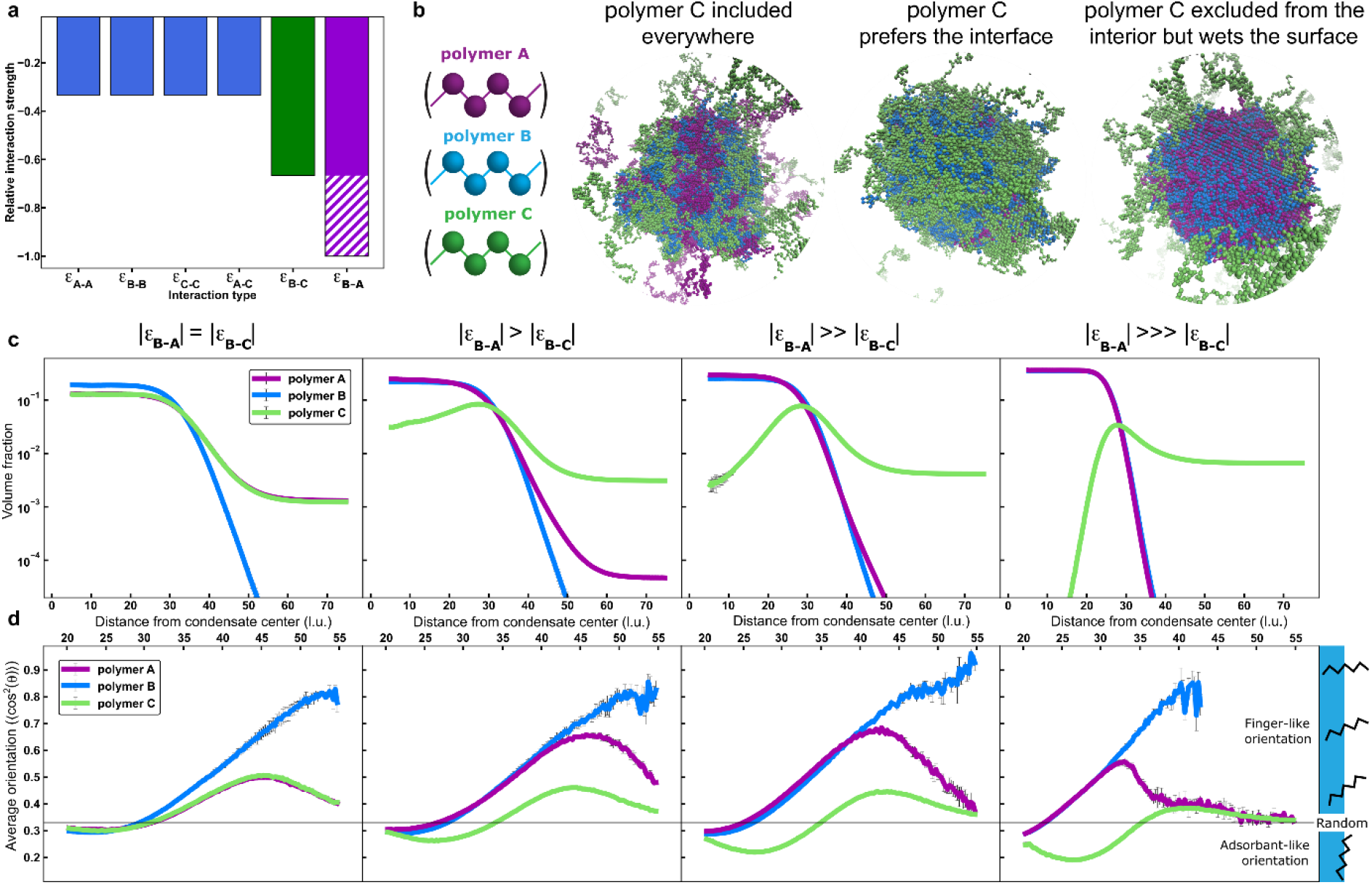
Pairwise interaction strengths and relative concentrations determine condensate architectures and interfacial features. **a**. The relative interaction strengths between polymer beads used in the simulations. The interaction between beads of polymers A and B was titrated from –0.67 to –1. **b**. Representative snapshots from LaSSI simulations showing distinct condensate architectures obtained by titrating the interaction strength between beads of polymers A and B. **c**. Radial density profiles of polymers A, B, and C in simulations with, from left to right, |ε_B-A_| = –0.67, –0.71, –0.75, and –1. **d**. Average cos^2^θ values, which indicate the average orientation of molecules with respect to the centre of the condensate for the corresponding systems described in **c** (Supplementary methods). The horizonal axis is rescaled from **c** to focus on the condensate interface. If a given datapoint had an error greater than 0.1, then that datapoint was omitted. In **c** and **d**, error bars indicate the standard errors about the mean across three replicates.

The microscopic pairwise interactions (ε_x-y_) between beads on polymers were chosen such that the three polymers engage in relatively weak interactions amongst themselves, except for B-A and B-C interactions, which are relatively strong with respect to thermal energy (**Figure 4a**, see Supplementary Methods). By maintaining equal polymer concentrations and titrating ε_B-A_, we obtained three possible architectures: (i) if |ε_B-A_| = |ε_B-C_|, then the polymers undergo uniform mixing within a single condensate; (ii) if |ε_B-A_| is slightly greater than |ε_B-C_|, then polymer C is partially displaced from the condensate interior to the condensate interface; and (iii) if |ε_B-A_| is significantly greater than |ε_B-C_|, then polymer C is completely excluded from the condensate interior, but can still preferentially interact with the interface of the condensate (**Figure 4b-c**). We confirmed that adsorption of a polymer requires for it to have a relatively weak interaction with the scaffolds that drive condensate formation. Additionally, we note that adsorption was realizable in scenarios (ii) and (iii).

Next, we titrated the concentrations of polymer A and polymer B, while keeping |ε_B-A_| slightly greater than |ε_B-C_| (scenario ii) (**Supplementary Figure 5**). Decreasing the amount of polymer B or increasing the amount of polymer A resulted in complete exclusion of polymer C from the condensate with little to no preference for the interface, suggesting that the polymer B is completely used up by polymer A. In contrast, increasing the amount of polymer B or decreasing the amount of polymer A resulted in a significant uptake of polymer C from the dilute phase into the condensate, thereby creating a core-shell architecture with a core composed of polymers A and B, and a shell composed of polymers B and C ^12, 23, 24, 25^. Finally, we titrated the concentration of polymer C while maintaining the interaction strengths as in **Figure 4d** and equal amounts of polymers A and B (**Supplementary Figure 6**). Regardless of whether the concentration of polymer C is less than or greater than that of polymers A and B, we see a clear preference for polymer C to adsorb onto the interface without the creation of a core-shell architecture. These simulations demonstrate how the interplay among polymer-polymer interaction strengths and relative concentrations can be used to fine-tune condensate organization.

### Adsorbents and scaffolds are oriented differently at the interfaces of condensates

Recently, Farag et al.,^26^ developed and deployed a specific order parameter aimed at quantifying the orientation of polymers with respect to the interfaces of condensates. In their simulations, the overall system had only one type of polymer *viz*., a coarse-grained, single bead per residue version of a prion-like low complexity domain. Phase separation in this system is driven by purely homotypic effects encoded by a hierarchy of sticker-and-spacer interactions. Farag et al., ^26^ found that polymers at the interface of the condensate were more likely to be oriented perpendicular to the interface. This minimizes the interfacial free energy density by maximizing the number of polymers at the interface and minimizing the number of un-crosslinked stickers per polymer.

Here, we employed a similar orientational analysis to determine whether the scaffolds and adsorbents adopt distinctive or similar orientations at the condensate interface. Specifically, we determined the angle θ swept out by the condensate centre-of-mass, a given polymer bead, *i*, and another bead on the same polymer that is exactly 100 beads apart from bead *i*. If θ is close to 0° or 180°, which corresponds to cos^2^θ values close to 1, then the 100-bead segment is perpendicular to the interface. In contrast, if θ is close to 90°, which corresponds to cos^2^θ values close to 0, then the 100-bead segment is essentially parallel to the interface. We refer to these limiting modes as being *finger-like* versus *adsorbent-like*.

**Figure 4d** shows the average orientations of polymers A, B and C under the same conditions as **Figure 4c**. We compare our results to a reference value of cos^2^θ = 0.33, which is the value we calculate for polymers with random orientations (**Supplementary Figure 7**). When |ε_B-A_| = |ε_B-C_|, all chains show a slight preference for finger-like orientations near the condensate interface, with the effect being most pronounced for polymer B. As |ε_B-A_| is increased to be above |ε_B-C_|, polymers A and B maintain their finger-like orientations near the condensate interface. Additionally, we observe a depletion of these species at the interface, and there is also significant dilution of these species in the coexisting dilute phase (**Figure 4c**). In contrast, polymer C shows an increasing preference for adsorbent-like orientations. These results suggest that polymers that behave as scaffolds, namely polymers A and B, minimize the interfacial free energy density by adopting finger-like orientations at the interface. In contrast, polymer C effectively becomes surfactant-like ^27^, in that the overall interfacial free energy density is minimized by adsorbent-like behaviour. The surfactant-like behaviour is not intrinsic because polymer C can form condensates on its own in the appropriate concentration regimes. Instead, the surfactant-like behaviour is an emergent property whereby condensate formation by the scaffold molecules causes adsorption if the relative interaction strengths are such that |ε_B-A_| > |ε_B-C_|. Interfacial adsorption maximizes interactions with the condensate interface while minimizing the disruption of favourable A-B interactions that provide the cohesive interactions for condensate formation and stabilization.

## Discussion

Our experiments and simulations help explain why certain macromolecules adsorb to the interfaces of condensates ^28^. Below its saturation concentration for phase separation, a macromolecule can be recruited to the interface of a condensate composed of scaffolds with lower saturation concentrations, and hence stronger driving forces for phase separation. At the interface of the PEG-poly-rA condensate, poly-rC molecules adsorb to drive a cascade of transitions from partial to complete wetting ^17^. When the wetting transition results in two coexisting dense phases, poly-rC adsorbs to the interface between the dense phases. This can be attributed to poly-rC taking advantage of the available PEG sites on the interface of the condensate.

Our observations, which were derived using generic, biocompatible, and biologically relevant polymers, help explain the phenomenology of adsorption that has been reported in cells ^17, 29^. Further, since multiphase condensates create internal interfaces ^10^, we propose that the internal interfaces between dense phases might provide additional spatial and temporal control over biochemical interactions. Specifically, interfaces between dense phases, as well as interfaces between coexisting dilute and dense phases, are likely to provide distinct molecular microenvironments for molecules to accumulate and interact. This is relevant given observations from the microdroplet literature, which show that reaction efficiencies can be enhanced by orders of magnitude at the interfaces of coexisting phases ^30^. In spatially organized condensates featuring *n* compositionally distinct phases, there are multiple internal interfaces that can be engineered to minimize off-target reactions in biochemical reactions involving multi-enzyme pathways ^30^.

In this work, we showed that a high concentration of macromolecules can accumulate at the periphery of condensates via different degrees of adsorption. Macromolecules defined by weaker driving forces for macroscopic phase separation are adsorbents recruited to interfaces of condensates formed by scaffold-like macromolecules defined by stronger driving forces for phase separation. Our computations show that scaffolds and adsorbents have different preferences for how they are oriented at the interfaces of condensates.

The mechanisms of adsorption and wetting are distinct from the site-specific effects of ligands that are known to modulate driving forces for phase separation through preferential binding across phase boundaries ^31, 32, 33, 34^. While preferential binding of ligands ^35^ can either compete against or add to extant interactions, thereby destabilizing or stabilizing condensates, adsorption of a ternary component has little effect on the interactions within a condensate ^36^. Overall, our results highlight key physical principles that govern the formation of spatially heterogeneous condensates of the type observed in many cells. Additional consideration of site-or chemistry-specific associative interactions are likely to provide further tunability to the spatial organization of macromolecules within and around condensates.

## Supporting information

Supplementary Information: Supplementary figures 1-7 and materials and method section

## Supplementary Information

Detailed description of the design of the experiments, reagents used, analysis pipelines and computational methods are available in the supplementary information document, as are additional supplementary figures.

## Acknowledgements

This work was funded by the Royall Scholarship (N.A.E.), the Center for Biomolecular Condensates at Washington University in St. Louis (N.A.E., R.V.P.), the European Union’s Horizon 2020 research and innovation programme under the Marie Skłodowska-Curie grant MicroREvolution (agreement no. 101023060; T.S.), the Winston Churchill Foundation of the United States (T.J.W.), the Harding Distinguished Postgraduate Scholar Programme (T.J.W.), Global Research Technologies Novo Nordisk A/S (H.A, T.P.J.K.), the US National Institutes of Health (R.V.P), the US Air Force Office of Scientific Research (R.V.P), the European Research Council under the European Union’s Seventh Framework Programme (FP7/2007-2013) through the ERC grants PhysProt (agreement no. 337969; T.P.J.K.) and the Newman Foundation (T.P.J.K., T.S.).

## Author Contributions

N.A.E., T.S. and T.P.J.K. conceived the study. N.A.E., M.F., D.Q., T.S., T.J.W., H.A., D.A.W., R.V.P. performed investigations and interpreted the results. T.P.J.K. and R.V.P. acquired funding. N.A.E., M.F., R.V.P. and T.P.J.K. wrote the original drafts, all authors reviewed and edited the paper.

## Additional information

Data generated during the study are available on reasonable request from the corresponding authors.

