## Supplementary Information: Supplementary figures 1-7 and materials and method section for "Adsorption of RNA to interfaces of biomolecular condensates enables wetting transitions"

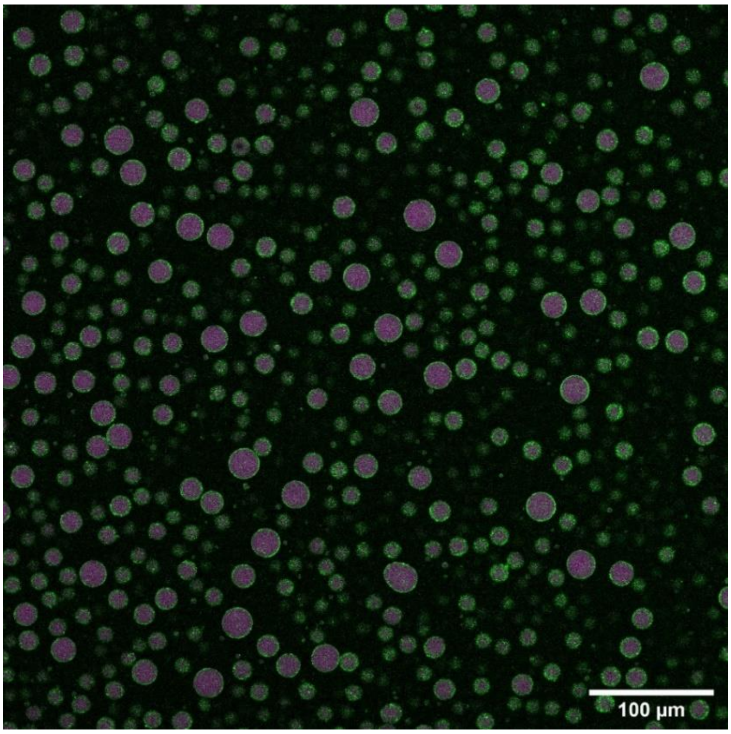


**Supplementary Figure 1: Overview confocal image of poly-rC (green) adsorbing to poly-rA (magenta) condensates.** Confocal image showing a large area of condensates with poly-rC adsorbing to the interface of Poly-rA enriched condensates. Samples contained 1000 ng/µL poly-rA, 1000 ng/µL poly-rC, 4 w/w% PEG, 750 mM NaCl and 50 mM HEPES at pH = 7.3.

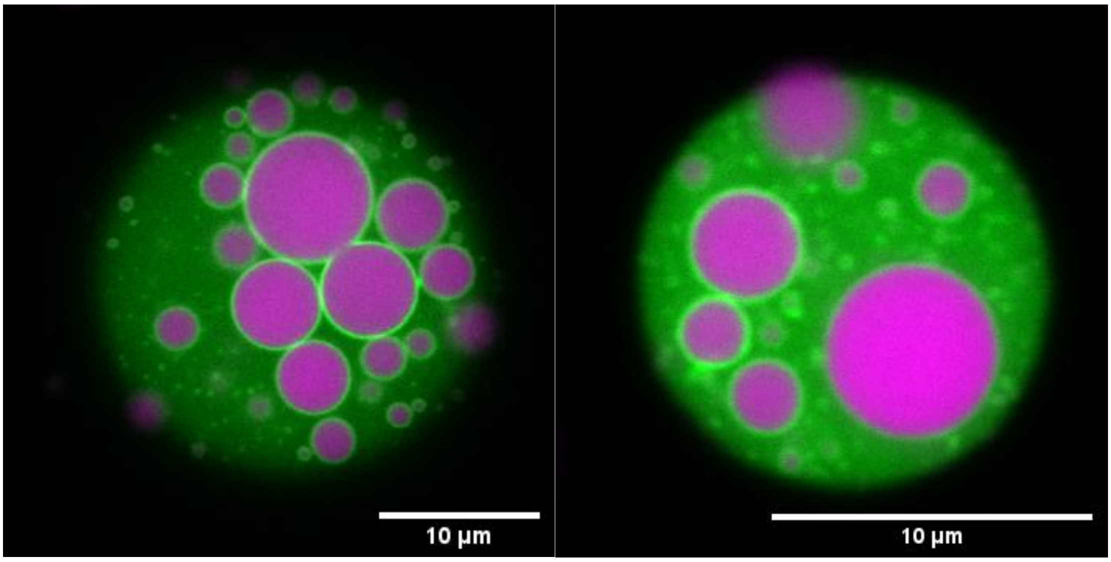


**Supplementary Figure 2: Additional confocal images of adsorption of poly-rC at the interface of coexisting dense phases.** Confocal images of multiphase condensates made of poly-rA (magenta), poly-rC (green) and PEG in aqueous solution. A poly-rC enriched phase surrounds a poly-rA enriched phase. Notably, the poly-rC adsorbs to the interface between the dense phases. The sample contained 1000 ng/µL poly-rA, 1000 ng/µL poly-rC, 8 w/w% PEG 20.000, 750 mM NaCl and 50 mM HEPES at pH = 7.3.

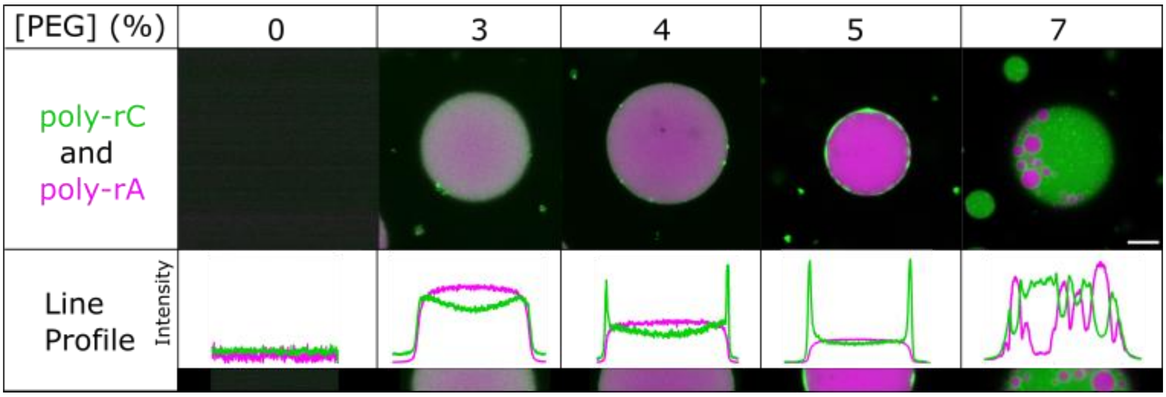


**Supplementary Figure 3: Line profiles of confocal images in figure 2d.** As the amount of PEG increases, poly-rC starts to absorb to the interface, partially wet and completely wet. Samples contain 1000 ng/µL poly-rA and/or poly-rC, a variable amount of PEG, 750 mM NaCl, and 50 mM HEPES pH = 7.3. The scale bar representing 10 µm applies to all confocal images.

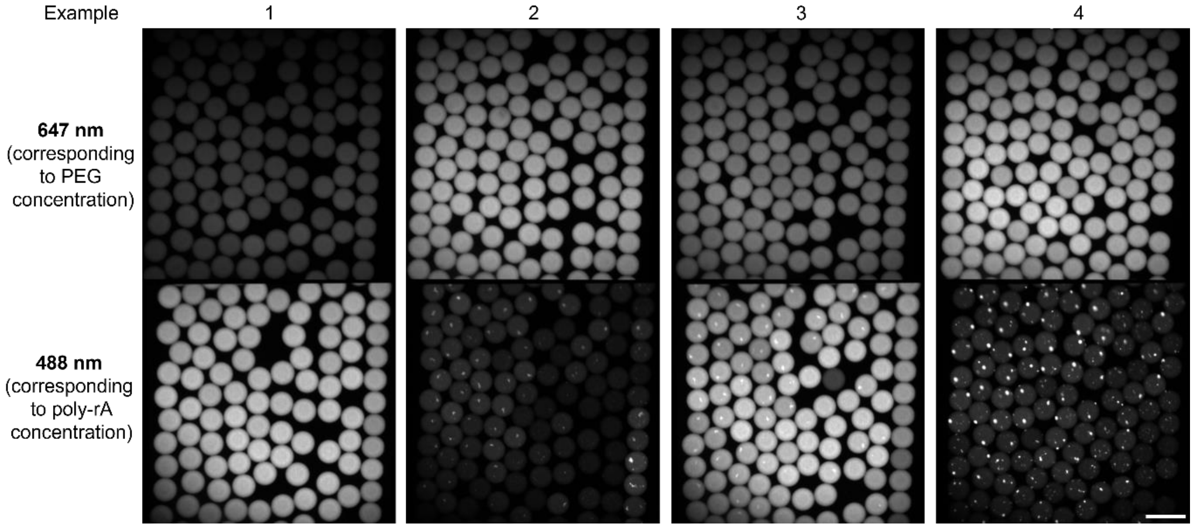


**Supplementary Figure 4: Examples of samples for Figure 3.** Wide-field fluorescent pictures of microfluidic droplets containing PEG, Alexa Fluor 647 carboxylic acid (concentration 0 – 10 µM relative to the PEG concentration), poly-rA (partly labelled with fluorescent dye at 488 nm), 750 mM NaCl, 50 mM HEPES pH = 7.3. Droplets are incubated for approximately 5 minutes before being imaged. Pictures are analysed by a python script adapted from [1]. When both a sufficient amount of PEG and poly-rA are present, condensates are formed. The 488 nm dye is attached to poly-rA and condensates are enriched in poly-rA. Thus, condensates can be seen as bright specs in droplets at 488 nm in example 2,3 and 4. From the intensity at 647 and 488 nm, we obtain the concentrations after comparing to the calibration curve. In this manner, we obtain both the total PEG and poly-rA concentrations and the dilute phase poly-rA concentrations. Combining this information with the information if these droplets contain condensates or not, we obtain the graphs in Figure 3. See the method and materials section for more details. The scale bar is 200 µm and applies to all images.

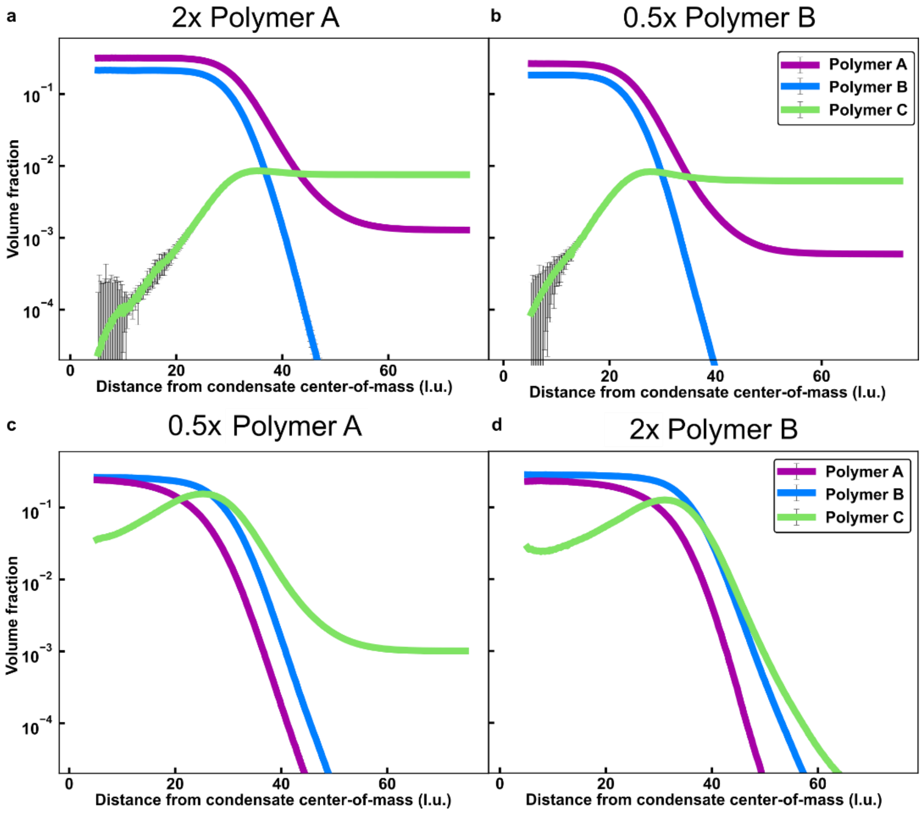


**Supplementary figure 5: Polymer A and B concentration titrations.** Radial density distributions of simulations with 100 molecules of polymer C and varying amounts of polymers A and B. **a.** 100 molecules of polymer A and 50 molecules of polymer B. **b.** 200 molecules of polymer A and 100 molecules of polymer B. **c.** 100 molecules of polymer A and 200 molecules of polymer B. **d.** 50 molecules of polymer A and 100 molecules of polymer B. The relative interaction energies are the same as those used in the third panel of Figure 4c.

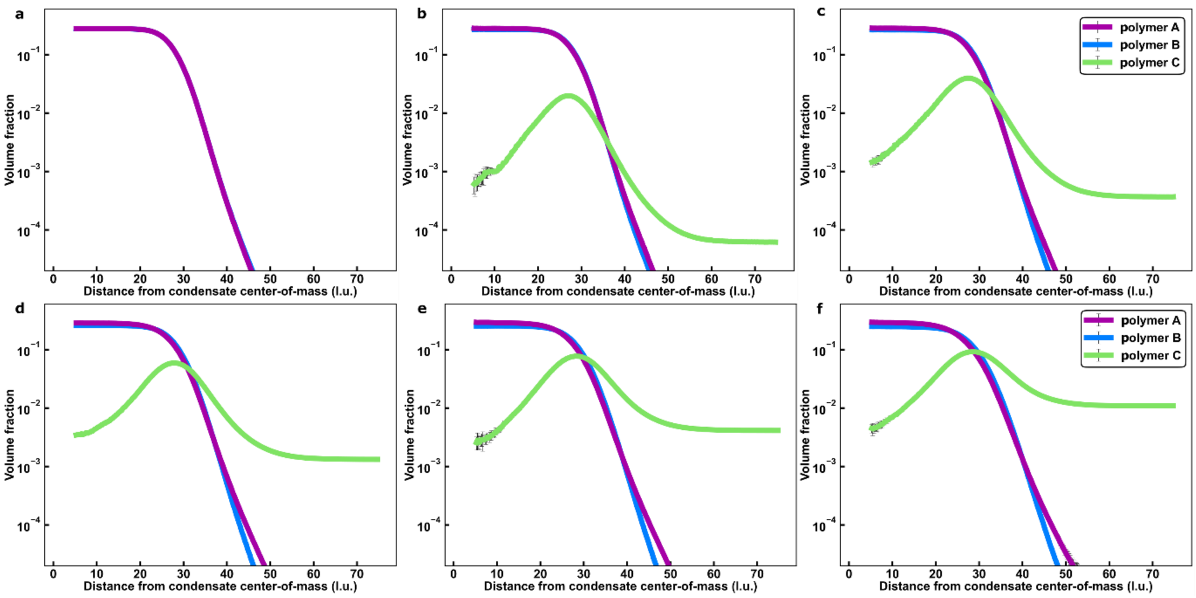


**Supplementary figure 6: Polymer C concentration titration.** Radial density distributions of simulations with 100 molecules of polymer A, 100 molecules of polymer B, and **a.** 0, **b.** 10, **c.** 25, **d.**, 50, **e.** 100, and **f.** 200 molecules of polymer C. The relative interaction energies are the same as those used in the third panel of Figure 4c.

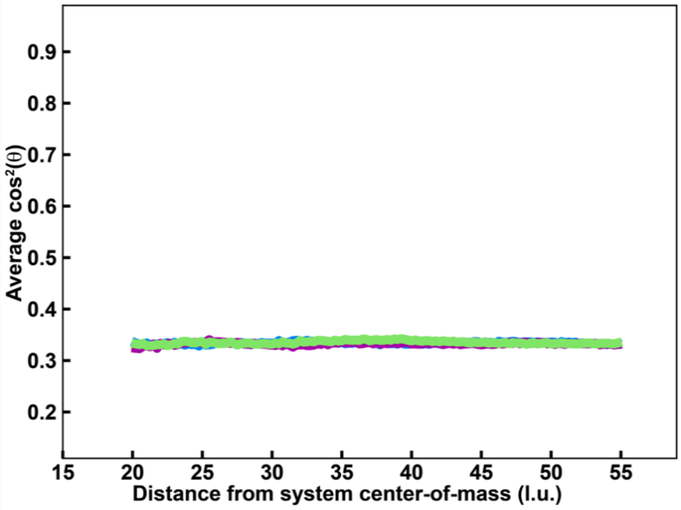


**Supplementary figure 7: Orientational analysis above the critical temperature.** Average $\cos^2\theta$ values (see Supplementary Methods) of polymers A, B, and C at a simulation temperature of 60. At this temperature, the system no longer undergoes phase separation, resulting in random molecular orientations. The average $\cos^2\theta$ value is about 0.33 at all distances for each polymer species. The relative interaction energy between polymer A and polymer B beads is 0.67. Simulations contain 100 molecules each of the polymers designated as A, B, and C. Standard errors about the mean across 3 replicates are smaller than the markers.

### Materials and Methods

### Materials

Poly-rA (MW 700-3500 kDa), PEG (average MW ~20,000), Poly-rC (average MW ~118 kDa), HEPES, NaCl, KCl, ethanol and mPEG(5k)silane were obtained from Sigma Aldrich. Alexa Fluor™ 647 Carboxylic Acid, Alexa Fluor™ 488 Carboxylic Acid and RNAase free water were obtained from Thermo Fisher. PEG(20k)-AF647 was purchased from Nanocs. Sylgard 184 Elastomer base and curing agent were purchased from Dow Corning Corporation. 18×18 mm glass slides were purchased from Academy. 24×60 mm No.1.5 glass slides were purchages from DWK Life Sciences. $(T)_{20}$-AF488 and $(C)_{20}$-AF546 were purchased from GenScript.

#### Fabrication of imaging wells for confocal microscopy

PDMS slabs of 3 mm in height were fabricated by mixing Sylgard 184 Elastomer base and curing agent at a 10:1 ratio and heating at 65 °C for 1.5 hours. Holes of 5 mm in diameter were punched in PDMS slabs of ~3 mm height using biopsy puncher. These slabs were plasma bonded to 24×60 mm No.1.5 cover glass slides to create wells. Both the wells and 18×18 mm glass slides used to seal the top of wells were treated with PEG-silane, using a method based on previously reports.[2,3] Briefly, the treatment solution was prepared by mixing 10 mg of PEG(5000)silane with 20 µL of glacial acetic acid and 1 mL of ethanol. Wells and glass slides were treated by placing them in solution for 1 hour at 65 °C and afterwards washing them thoroughly with water. Treated devices are not used after more than 3 weeks since the treatment.

### Condensate preparation

Condensates of poly-rA and or poly-rC with PEG were prepared by mixing stock solutions in an Eppendorf tube in the following order: buffer, salt, PEG, pre-stained poly-rC and / or pre-stained poly-rA. All solutions were prepared in RNAase free water. The poly-rA and poly-rC concentrations were determined with a nanodrop machine by measuring the absorbance at 260 nm. Poly-rA and poly-rC were fluorescently labelled by incubation of 0.5 mol-% of $(T)_{20}$-AF488 or $(C)_{20}$-AF546 for 3 minutes at 80 °C and excess dye was removed using an RNA clean up kit (Qiagen). The sample was placed in an imaging well and sealed to be air tight.

**Confocal imaging**

A Leica Stellaris 5 confocal microscope (confocal fluorescence imaging) (white light laser) microscope equipped with a 63× oil immersion Leica 1.4 NA was used for imaging. Further analysis of pictures was performed using Fiji. In **Figures 1 and 2**, the legend applies to all images to the left until either a new legend is encountered or the end of the image. Different legends were made so that the intensity can be compared, and structures remain easily visible.

**Microfluidic device design and fabrication**

Standard lithography processes were used to fabricate the microfluidic devices.[1,4] AutoCAD (AutoDesk) software was used to design the devices, which were printed on a photomask (Micro Lithography). A 45 µm layer of SU8-3050 photoresist (Microchem) was spun on a silicon wafer and heated at 95 °C for 15 minutes. The photomask was placed on top, and the pattern was placed onto the wafer using exposure to UV light and heating at 95 °C for 15 minutes. Propylene glycol methyl ether acetate (PGMEA) was used to remove excess unreacted SU8-3050. The wafer was dried and a profilometer was used to confirm the height of the device. PDMS at 1:10 curing agent:base ratio was placed on top of the mould and heated at 65 °C for 1.5 hours. The PDMS was cleaned, holes were punched into inlets and outlets, the PDMS substrate was then bonded to glass slides using a plasma oven (30 s, 40% power, Femto, Diener Electronics). The interior of the device was treated with 1% trichloro(1H,1H,2H,2H-perfluorooctyl)silane (Sigma) in HFE-7500 (Fluorochem), before being dried and heated at 95°C for 1 min.

**Mapping phase diagrams and constructing tie-lines**

Aqueous droplets in oil of an average volume of 100 picolitre were created at approximately 100 droplets per second using a microfluidics setup as shown schematically in **Figure 3a** of the main text. Specifically, a solution of fluorescently labelled $10^4$ ng/µL poly-rA or poly-rC was mixed with a solution of fluorescently labelled 20 or 10 w/w % PEG and a solution with buffer at varying rates, while being encapsulated in fluorinated oil (HFE-7500 Fluorochem with 1 v/v % fluorosurfactant). To flow the solutions into the microfluidic device, syringes (Hamilton 1710) and syringe pumps (neMESYS modules, Cetoni) were used. Syringes were connected to the microfluidic device using tubing (PTFE, 0.012"ID × 0.030"OD, Cole-Parmer). The 100 picolitre droplets were created by a total aqueous flow rate of 60 µL/h and the oil flow rate of 150 µL/h. After a 5-minute incubation period, the aqueous droplets in oil were imaged simultaneously at the wavelength corresponding to poly-rX (488 or 546 nm) and PEG (647 nm). These images were analysed using the Python program[1] to determine the concentrations of these compounds and the presence or absence of condensates. Additionally, the dilute phase concentration of poly-rX was determined by calibrating the intensity of 488 or 546 in an area absent of condensates to concentration poly-rX concentration (**Figure 3b**). Each droplet gives information for one datapoint in **Figures 3c or 3d**. The boundaries were determined using fitting that is based on support vector machine-based methods. The tie-lines were fitted between the "condensates" datapoints by grouping them based on their poly-rX dilute phase concentration. **Figures 3c and 3d** show the datasets and the tie-line when grouping the data into 0 – 500, 500 – 1000 and above 1000 ng/μL. Notably, to determine slopes and their errors, the data were grouped $10^3$ times with different cut-offs. Using these $10^3$ angles, we determined the average and standard deviation. This way, we did not have an arbitrary cut-off and were able to determine the standard deviation of the measurement method.

#### Coarse-grained simulations

Simulations were performed using LaSSI, a lattice-based Monte Carlo engine[5]. Monte Carlo moves are accepted or rejected based on the Metropolis-Hastings criterion so that the probability of accepting a move is equal to min(1, exp(–$\beta\Delta E$)), where $\beta = 1 / kT$. Here, $kT$ is the simulation temperature (set to 50) and $\Delta E$ is the change in total system energy associated with the attempted move. Total system energies were calculated using a nearest neighbour model and the following interaction energies: $e_{A\text{-}A} = \varepsilon_{B\text{-}B} = \varepsilon_{C\text{-}C} = \varepsilon_{A\text{-}C} = -0.33$; $\varepsilon_{B\text{-}C} = -0.67$; $\varepsilon_{B\text{-}A}$ was –0.67, –0.71, –0.75, or –1. Unless stated otherwise, each simulation involves $10^2$ distinct polymers of each type comprised of 250 beads each in a cubic lattice with length 150 lattice units. The simulations were performed for $5\times10^{10}$ Monte Carlo moves and were allowed to equilibrate such that the chains formed a single condensate with a coexisting dilute phase before any analysis was performed. Three independent simulations were performed at each condition, and the results shown are aggregates over all replicates. Radial densities of each polymer type were normalized using the exact number of lattice sites in each radial shell. The condensate centre-of-mass was taken to be the centre-of-mass of the single largest network of interacting chains. The chain orientation analysis in **Figure 4d** was introduced previously[6]. Here, we perform the analysis as follows: (1) for a given chain in the system, find all pairs of beads ($i$, $j$) that are exactly 100 beads apart along the chain. (2) For each pair, draw a line segment from the condensate centre-of-mass to bead $i$. (3) Draw a line segment from bead $i$ to bead $j$. (4) Determine the angle, $q$, swept out by the two different line segments. (5) Calculate $\cos^2\theta$ and bin this value based on the distance between bead $i$ and the condensate centre-of-mass. (6) Repeat steps 1-5 for all chains in the system. (7) For each bin (as described in step 5), we report the average of the calculated $\cos^2\theta$ values. If a given datapoint had an error greater than 0.1, then that datapoint was omitted.
